# Guardians of our Rivers: transforming river health assessment through macroinvertebrate monitoring with citizen scientists

**DOI:** 10.64898/2026.08.26.739408

**Authors:** Rebecca Lewis, Kerry Dodd, Craig Macadam, Iain Matthews, Stefan R. Pulver

## Abstract

Freshwater ecosystems in Scotland are increasingly threatened by multiple interacting stressors, including pollution, hydrological alteration, land use change, and climate-driven pressures. Scotland’s rivers are amongst the most physically diverse and dynamic in the UK, contributing significantly to the country’s economy and natural heritage. Monitoring 125,000 km of waterways presents an issue, with limited funding and capacity of environmental agencies creating an environmental monitoring deficit at a time when robust data are essential for delivering national and global biodiversity targets. Citizen science offers a scalable, community driven approach to strengthening environmental evidence. This study evaluates the development, implementation, and outcomes of *Guardians of our Rivers* (GooR), a nationwide citizen science programme using macroinvertebrate-based river health assessments based on the Anglers’ Riverfly Monitoring Initiative (RMI) established by the Riverfly Partnership. Between 2022 and 2025, GooR trained more than 850 volunteers, established 123 monitoring sites across Scotland, and generated over 860 surveys—surpassing the previous 16 years of aquatic invertebrate monitoring efforts. Standardised field training, full equipment provision, and structured support enabled high-quality data collection across diverse catchments. The establishment of nationally applied trigger and target thresholds created a consistent mechanism for detecting ecological deterioration and ensured that verified trigger breaches (n = 39 in 2025) resulted in formal investigation or follow-up. Analysis of national abundance patterns revealed ecological and biogeographic trends across the eight RMI taxa, confirming data sensitivity to habitat differences, water quality, and regional pressures. A detailed case study of the Lothian Esk catchment demonstrated the programme’s capacity to detect seasonal macroinvertebrate dynamics and site level ecological trajectories over time. Overall, GooR illustrates how well-designed and well-supported citizen science can deliver high resolution spatiotemporal datasets, enhance early warning systems, empower communities, and make a meaningful contribution to promoting river health. Importantly, GooR forged a working partnership with the Scottish Environmental Protection Agency (SEPA) that allowed communities to not only gather data but also transformed the way citizen science is recognized as contributing to freshwater conservation. Continued investment and long-term support will be essential to sustain these gains and realise the full potential of citizen-driven freshwater stewardship.

**Author Summary:** In this study, we explore how communities across Scotland can help protect the health of their rivers at a time when freshwater ecosystems are under increasing pressure from a range of threats. Scotland has an extensive network of rivers and streams, creating a challenge for environment agencies to provide comprehensive monitoring coverage that delivers the level of preventative care needed to keep our water courses in optimal health. To address this, we created a national citizen science monitoring programme, ‘*Guardians of our Rivers*’, and showed that by harnessing local knowledge and with training and support, local communities can collect robust reliable data about river health, contribute meaningfully to local environmental stewardship, and successfully engage with the relevant statutory body, SEPA.

Between 2022 and 2025 more than 850 volunteers learned how to assess river health by counting freshwater macroinvertebrates that are known to be sensitive to changes in water quality. Over 860 surveys were carried out at 123 monitoring sites, creating a level of coverage not possible with professional monitoring alone. Working together with SEPA, we established protocols to increase robustness and consistency in data collection and empower citizen scientists to share information, which can form the basis for further actions. Our work demonstrates how citizen science can produce robust data, strengthen existing monitoring efforts, and build stronger connections between communities and their rivers. Our works highlights a scalable model that could be implemented widely in the UK and beyond.

## Introduction

Freshwater ecosystems are facing unprecedented pressure both locally and globally (1) (2). Declines in the taxonomic and functional biodiversity of freshwater communities including microorganisms, algae, plants, invertebrates and fish can alter ecological processes and associated benefits to people (3). River habitats and their ecosystems are threatened by a range of stressors that are often highly interconnected: ongoing human development (4), (5), including the modification of channel morphology (6), dredging (7), changes to catchment land-use (8), pollution from diffuse and point sources(9, 10), invasion by alien species(11), and alterations of the flow regime from abstraction, coincide with damming and flood risk management (12). Since 1950, existing river stressors have intensified and new stressors have emerged, including neonicotinoid pesticides, veterinary flea products (13, 14), microplastics(15), pharmaceuticals and personal-care products (14) (16), worsening climate change (17) (18) and the development of global hydropower (19, 20). New stressors often co-occur with historical persisting stressors, potentially exacerbating their combined effects on river biodiversity (21, 22). These trends also have economic impacts; recent work suggests nature degradation could reduce UK GDP by around 12% (£150–300 billion) by 2030 (23). At the same time, the finance gap required to meet the UK’s nature related outcomes is estimated at £56 billion per year (23). Reversing the decline in freshwater health requires robust environmental monitoring data; however, shrinking core funding and capacity have created a critical information gap.

Faced with these challenges, river management bodies and government agencies need to prioritise restoration strategies that build ecological resilience, enabling ecosystems to adjust to future change while sustaining essential natural functions (24, 25, 26). Delivering ecosystem resilience requires actions with the right balance of complexity and scale to be both effective and sustainable (27). Balancing the demands of ensuring a legacy of healthy freshwater ecosystems for future generations with constraints of an information and financial deficit requires us to look for alternative collaborative solutions to achieve this resilience. Citizen science has become one of the most effective approaches for addressing environmental decline and supporting ecological recovery. It enables the collection of large quantities of data at scale, increasing both temporal and spatial resolution, while also building an informed and engaged community that supports sustainable environmental management and contributes to climate change mitigation (28, 29). Citizen generated data are now routinely integrated into delivery projects across governmental and nongovernmental organisations (30, 31, 32). Globally, interest in citizen science continues to grow, with platforms such as citizenscience.eu, SciStarter, and Zooniverse providing access to projects, resources, and training. Earthwatch alone supports around 1,400 field research projects in more than 120 countries, contributing over 10 million hours of data collection (33). Within the UK, citizen scientists working with the Riverfly Partnership—whose methodology underpins this research— monitor more than 4,600 sample sites and have collected over 71,600 samples (34). Creating resilient citizen science programmes is therefore essential to achieving these national and local goals.

Freshwater invertebrates have long been recognised as a powerful bioindicator of the condition of our freshwater habitats, with biological monitoring protocols used systematically for determining the ecological condition of water bodies since the development of the saprobic system, more than 100 years ago (35). Research on specific taxa at species, genus or family level has established tolerance limits to a range of environmental conditions, including salinity (36), pH (37), organic pollution (38), suspended sediment concentration (39), fine sediment deposition and flow velocity (40, 41). As many freshwater invertebrate taxa have narrow tolerance ranges to key abiotic conditions, they provide a sensitive tool for assessing both current health of these habitats and the effectiveness of interventions to improve and repair them (42). Selecting a suite of freshwater macroinvertebrates (also known colloquially as ‘Riverflies’) for use in citizen science projects not only provides a practical method for river health assessment but can also help develop public understanding of the role that invertebrates play in a healthy, sustainable trophic system (42).

Scotland contains more than 125,000 km of rivers and streams, ranging from small upland burns to large lowland systems such as the Tay, and these waters underpin national wellbeing, industry, recreation, and natural heritage (43). The Water Framework Directive (WFD) provides the legislative basis for protecting these ecosystems, placing strong emphasis on biological quality elements, including macroinvertebrates, fish, diatoms, and macrophytes, for assessing ecological status (44). Under the WFD, Scotland’s current River Basin Management Plans set the target for 81% of water bodies to reach “good” or better status by 2027 (45). Achieving this requires extensive, high-quality monitoring, yet the scale and remoteness of Scotland’s rivers, combined with limited institutional capacity, mean that statutory agencies may struggle to meet this demand alone. Addressing these gaps and improving river health therefore depends on wider community involvement and collaborative monitoring approaches.

Citizen science monitoring of freshwater ecosystems in Scotland has developed progressively over the past two decades, driven largely by the need to supplement limited statutory monitoring capacity. Early initiatives were small and locally organised, with volunteer groups undertaking ad hoc macroinvertebrate surveys to support river restoration projects and local biodiversity action plans. A major step change occurred in 2013 with the launch of the Clyde Riverfly Monitoring Partnership (CRIMP), which expanded RMI into Scotland, establishing long-term community led monitoring across the Clyde catchment. Between 2013 and 2022, additional Riverfly groups emerged across the central belt and the Borders, gradually expanding geographic coverage. However, expansion remained uneven, focussed around urban centres, and dependent on local champions. By the end of 2021, Scotland had 79 registered Riverfly monitoring sites and 22 volunteer groups, yet data entry had slowed down with only 77 surveys entered that year by only 5 remaining active groups. Large areas of the country—outside of the central belt —remained unmonitored. This historical foundation provided the platform upon which Guardians of our Rivers (GooR) was built, enabling a transition from fragmented local efforts to a coordinated, nationwide citizen science network.

The programme reported in this study undertook a nationwide engagement effort, training and supporting communities across Scotland to monitor the health of their local rivers using freshwater macroinvertebrates. The aim was to create resilient, community-driven citizen science groups that become stewards of their local rivers. A central objective was to establish a durable partnership with the Scottish Environment Protection Agency (SEPA) in a bid to facilitate the transformational change needed to protect our freshwater ecosystems. Without this collaboration, the full potential of citizen science for our rivers could not be fully realized. This study therefore seeks to 1) Describe the process of establishing GooR; 2) Document the growth of GooR and the expansion of monitoring sites; 3) Examine the historical data and contribution of GooR across Scotland; 4) Present a case study from an individual river catchment to illustrate ecological trajectories over time; and 5) Assess the implementation and value of site-specific triggers and targets levels established in cooperation with the relevant statutory body.

## Results

### Overview of Riverfly Monitoring data across Scotland

To understand the context for creating GooR, we first analysed historical Riverfly Partnership data available for Scotland. Beginning in 2009, citizen scientist groups in Scotland sampled macroinvertebrate diversity using the RMI method, developed as part of the Riverfly Partnership (46). The method involves standardized kick sampling (Figure 1A) followed by live sorting (Figure 1B) into 8 taxa, strategically chosen based on varied tolerances to pollution, distribution across UK, and ease of identification. The RMI indicator taxa are: Cased Caddis (Trichoptera); Caseless Caddis (Trichoptera); Mayfly (Ephemeroptera); Blue-winged Olive (Ephemerellidae); Flat-bodied (Heptageniidae); Olives (Baetidae); Stoneflies (Plecoptera); Freshwater Shrimp (*Gammarus* spp.). (Figure 1C) (see Materials and Methods for details).

**Figure 1.**
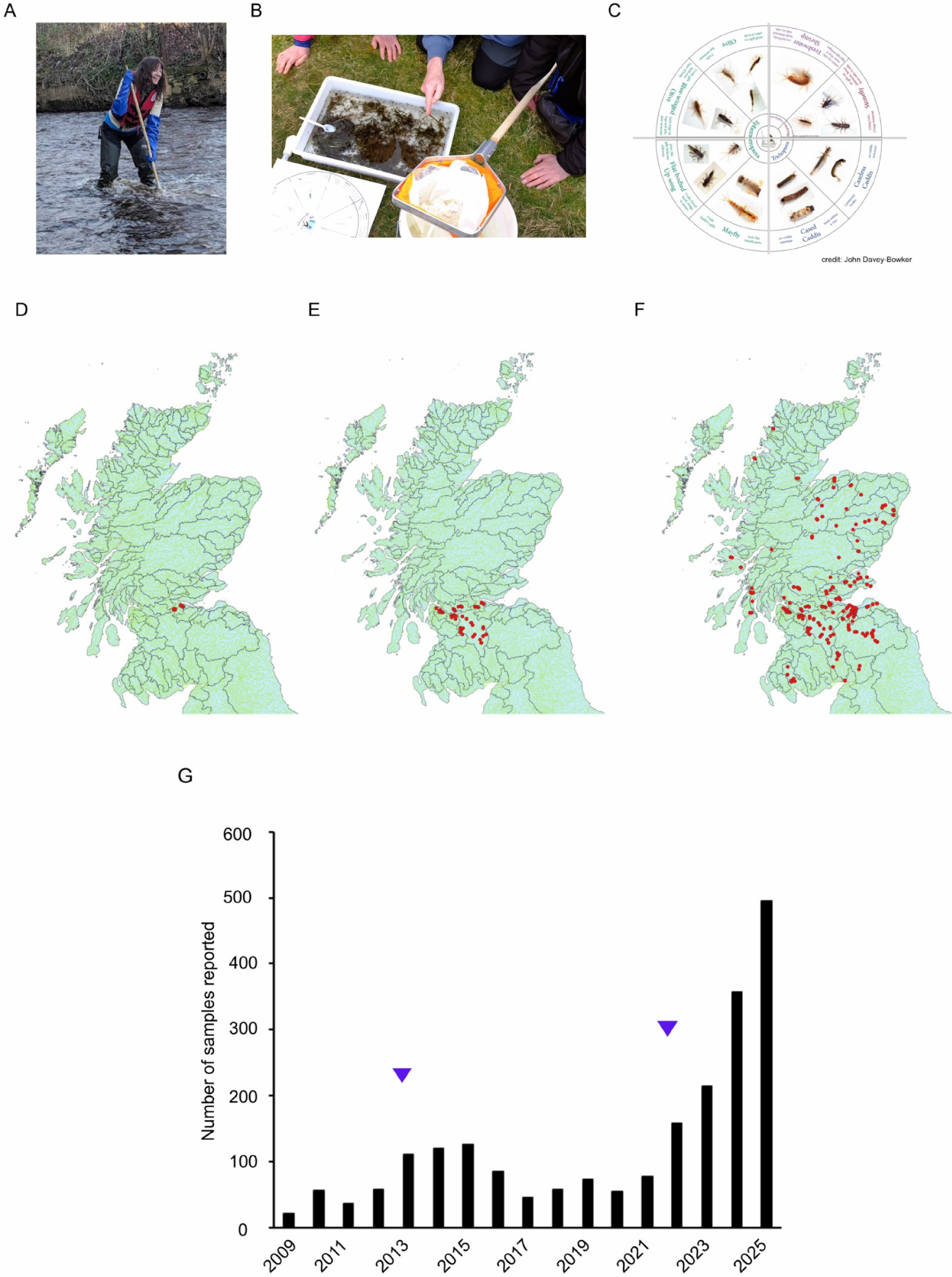
Overview of GooR activities and development over time. (A) Author Rebecca Lewis demonstrating kick-sampling methods to collect aquatic macroinvertebrates in rivers (RL consents to publication of photo); (B) GooR group live sorting collected samples; (C) images of 8 RMI taxa; (D) Sites at which RMI sampling data were recorded 2009, 2013(E), 2025(F). (G) Complete history of RMI sampling data collected in Scotland since 2009. Blue arrowheads indicate date of initiation of Riverfly monitoring projects (CRIMP, left) and GooR (right)

A set of initial sites were established in 2009 (Figure 1D); in 2013, the CRIMP project along the River Clyde was launched, resulting in an increase in sites (Figure 1E); in 2018 – 2022 monitoring sites spread across several river catchments in the central belt and south into the Scottish Borders. In late 2022, Buglife Scotland launched GooR. Buglife promoted GooR in a media campaign, then coordinated and delivered training sessions across multiple sites in Scotland ahead of the 2023 field season (see Materials and Methods for details). Subsequently, in 2023 – 2025, there was a wide-scale spread of monitoring sites across Scotland, including the West and East coasts, as well as Highlands and Islands areas (Figure 1F). At the end of 2025, 123 monitoring site locations were registered in the GooR project across Scotland. The number of monitoring locations per river catchment ranged from 1-15. A total of 1,237 surveys were entered onto the Riverfly database between 2009 and 2025 from 22 original Riverfly groups encompassing 97 sites in Scotland. Since GooR officially launched in October 2022 until December 2025, 861 surveys were entered into the database by 58 additional groups, encompassing at total of 123 monitoring sites (Figure 1G). The number of volunteers trained increased over each of the three GooR years from 99 in 2023, to 304 in 2024 to 458 in 2025. A total of 859 volunteers were trained over 2023-2025 in the GooR program.

**Figure 2.**
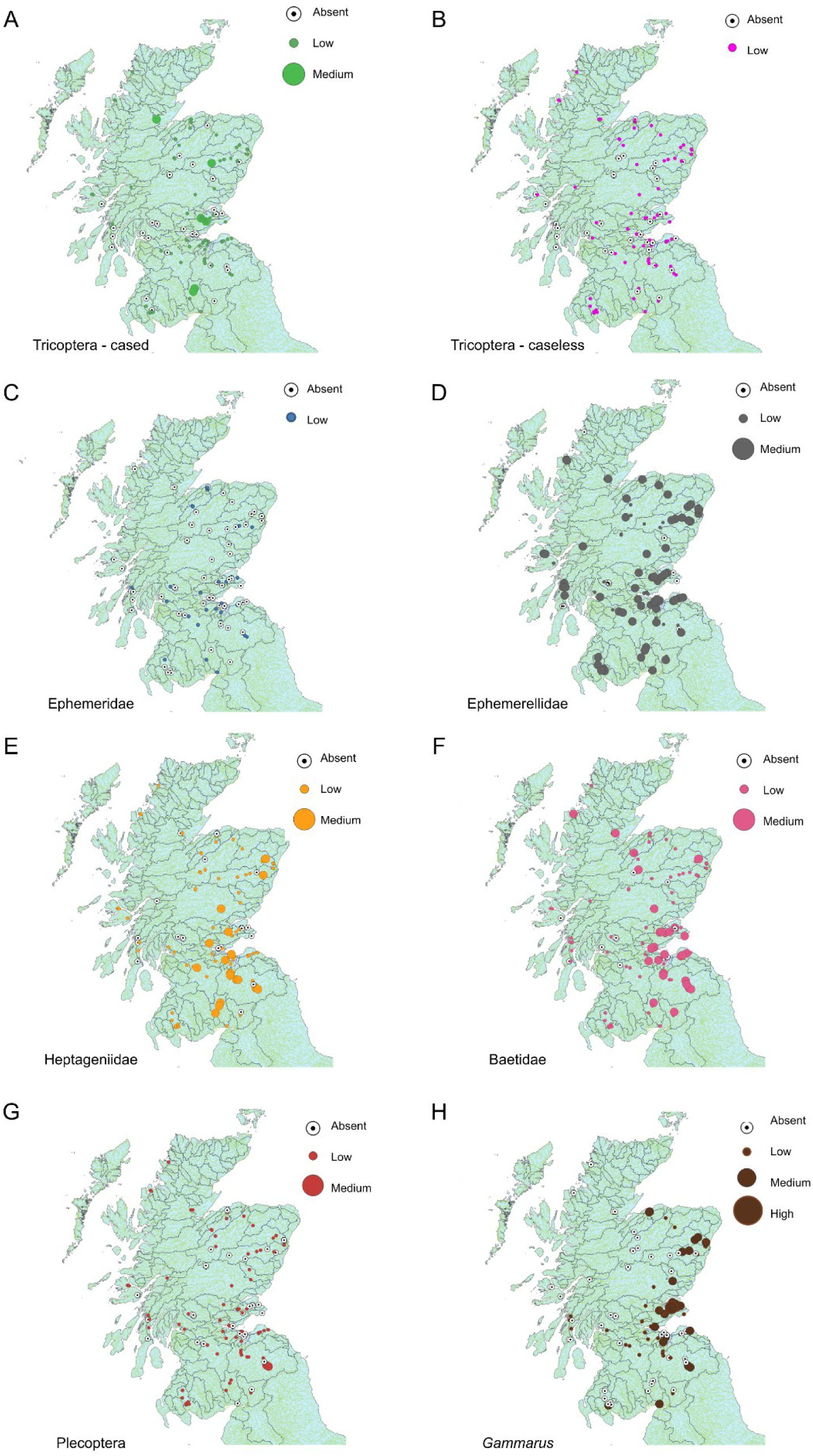
Geographic spread and abundance of each of the RMI 8 invertebrate groups in each of the sampling sites across Scotland. (A) Trichoptera - cased, (B) Trichoptera - caseless, (C) Ephemeridae, (D) Ephemerellidae, (E) Heptageniidae, (F) Baetidae, (G) Plecoptera, and (H) *Gammarus* mean score calculated and categorised into Absent, Low, Medium and High abundances.

RMI indicator species were originally chosen in part because they are widely distributed across England, and easily identified; however, few studies to date have systematically examined the geographical distribution of these groups across Scotland. This represented an opportunity for GooR to generate novel data. We therefore plotted the spatial distributions of RMI scores for each indicator taxa using the free open-source spatial mapping software QGIS (Figure 2). To allow for comparison across groups, we calculated mean RMI scores across all years of GooR, then categorized mean scores into *‘Absent’*, ‘*Low’*, ‘*Medium*’, and ‘*High’*. Across the surveyed sites, the abundance of Riverfly taxa was generally recorded in the *Low* (i.e. mean RMI score < 2) category, with most measurements dominated by *Absent* or *Low* RMI scores. Cased Caddis were largely present in the *Low* category (64%), with fewer *Absent* records (31%) and only a small proportion in *Medium* (i.e. mean RMI score = 2-3) category (5%). Caseless Caddis were restricted to *Absent* (34%) and *Low* (66%) categories, with no sites registering in *Medium* or *High* categories. Observations of Ephemeridae fell largely into the *Absent* category (71%) and the remainder in the *Low* category (29%). In contrast, Ephemerellidae displayed a stronger presence, with *Medium* forming the dominant category (59%), followed by *Low* (30%) and a smaller proportion absent (11%). Heptageniidae were recorded most frequently at *Low* category (60%), with additional *Medium* (17%) and *Absent* (23%) occurrences. Baetidae also showed a predominance of *Low* levels (63%), alongside *Medium* (26%) and a minority *Absent* (12%). *Gammarus* exhibited the broadest distribution across categories, with absent (44%) and ‘Low’ (35%) most common, followed by *Medium* (19%) and rarely in the *High* (i.e. mean RMI score 3-4) category (2%). Plecoptera were present mainly in the *Low* category (65%), with a third of records absent (33%) and only a small proportion in the *Medium* category (2%). Collectively, these patterns indicate that most taxa occurred primarily at *Absent* or *Low* categories with only a few groups achieving *Medium* or *High* abundance designations.

RMI taxa were, in most cases, widely distributed across sampling sites in Scotland. Caseless Caddis, Ephemerellidae and Plecoptera all showed fairly even distributions of abundance categories across monitoring sites. For Cased Caddis, small clusters of *Medium* levels appeared in a few isolated catchments, rather than showing any systematic, large scale geographical biases. Similar results were obtained with Heptageniidae and Baetidae, with Baetidae showing clusters of *Medium* abundance categories around the mouth of the Forth estuary on the east coast of Scotland. *Gammarus* were the only group showing some signs of geographical bias with a large number of *Medium* and *High* categories on the east coast compared to mostly *Absent,* or *Low* designations at west coast sites. Finally, Ephemeridae were largely *Absent* or extremely rare at most sites across Scotland.

### Case study: Long term seasonal trends on the Lothian Esk

In some instances, monitoring groups measure multiple sites within a given river catchment over time to examine long-term sampling trajectories, season trends, and site-specific conditions. One example is the ‘Riverfly-on-the-Esk’ group, which monitors the Lothian Esk, a river that flows through the Mid and East Lothian region of Scotland (Figure 3A). As of 2025, the Riverfly-on-the-Esk group had over 50 trained volunteers and 15 registered monitoring sites (Figure 3B). The group has been collecting data since 2019.

To examine how the abundances of RMI taxa varied across seasons in the Lothian Esk catchment, we plotted mean +/-95% confidence intervals for all data collected in the catchment each month (Figure 3C-J). Heptageniidae and Baetidae, were continually found in relatively high numbers, with monthly averages frequently reaching over 200 samples, and outliers reaching up to 600 for Heptageniidae, and over 350 for Baetidae. Despite these high peaks, mean values for both groups typically remained at or below 100. Average abundances of Cased caddis, Caseless caddis and Plecoptera typically fell in the range of 45-50, with outliers reaching 100 and over. Mean values for all three groups remained at or below 25. Most *Gammarus* abundances measurements fell below 100, with occasional high outliers exceeding 150. Mean monthly values did not exceed 25. Strikingly, Ephemeridae abundance remained consistently very low throughout the year, with most observations registering at, or very near zero individuals across all months with occasional higher counts reaching at most only 1 or 2 individuals.

Looking across seasons, Cased caddis showed variability in the data and no obvious seasonal pattern, small dips in the mean number were recorded in February and June. Caseless caddis also showed variability in the data but with a small rise in numbers from late spring to mid-summer (Figure 3C, 3D). Ephemeridae abundance remained consistently low across all months, with most observations at or near zero. Only infrequent counts of one or two individuals were recorded, and these did not follow any clear seasonal pattern (Figure 3E). Ephemerellidae abundance showed a pronounced seasonal pattern, with extremely low or zero abundances recorded throughout most of the year and a distinct surge in mid-spring (Figure 3F). Heptageniidae showed variability across months, but also a seasonal pattern, with a rise in mean numbers from late winter through to spring, dropping in summer and into early winter (Figure 3G). Baetidae showed substantial variability across months, but with a rise from late winter through to autumn and a slight drop in late autumn through to winter (Figure 3H). Plecoptera showed variability in the data but with relatively little seasonal variation (Figure 3I). Finally, the seasonal distribution of *Gammarus* abundances showed variability across months; typical monthly mean values did not fluctuate dramatically despite the presence of sporadic, high counts. The dispersion of data points increased from July to October, reflecting a wider range of abundances during warmer months, including the highest recorded values in the dataset. In contrast, winter and early spring months showed tighter clustering of observations, suggesting more stable and generally lower *Gammarus* populations during these periods.

Overall, seasonal abundance trends on the Lothian Esk were qualitatively consistent with averaged annual patterns seen in other regions of Scotland (Figure 2). In particular, data from the Lothian Esk were representative of the trend towards very few observations of Ephemeridae at GooR monitoring sites

**Figure 3.**
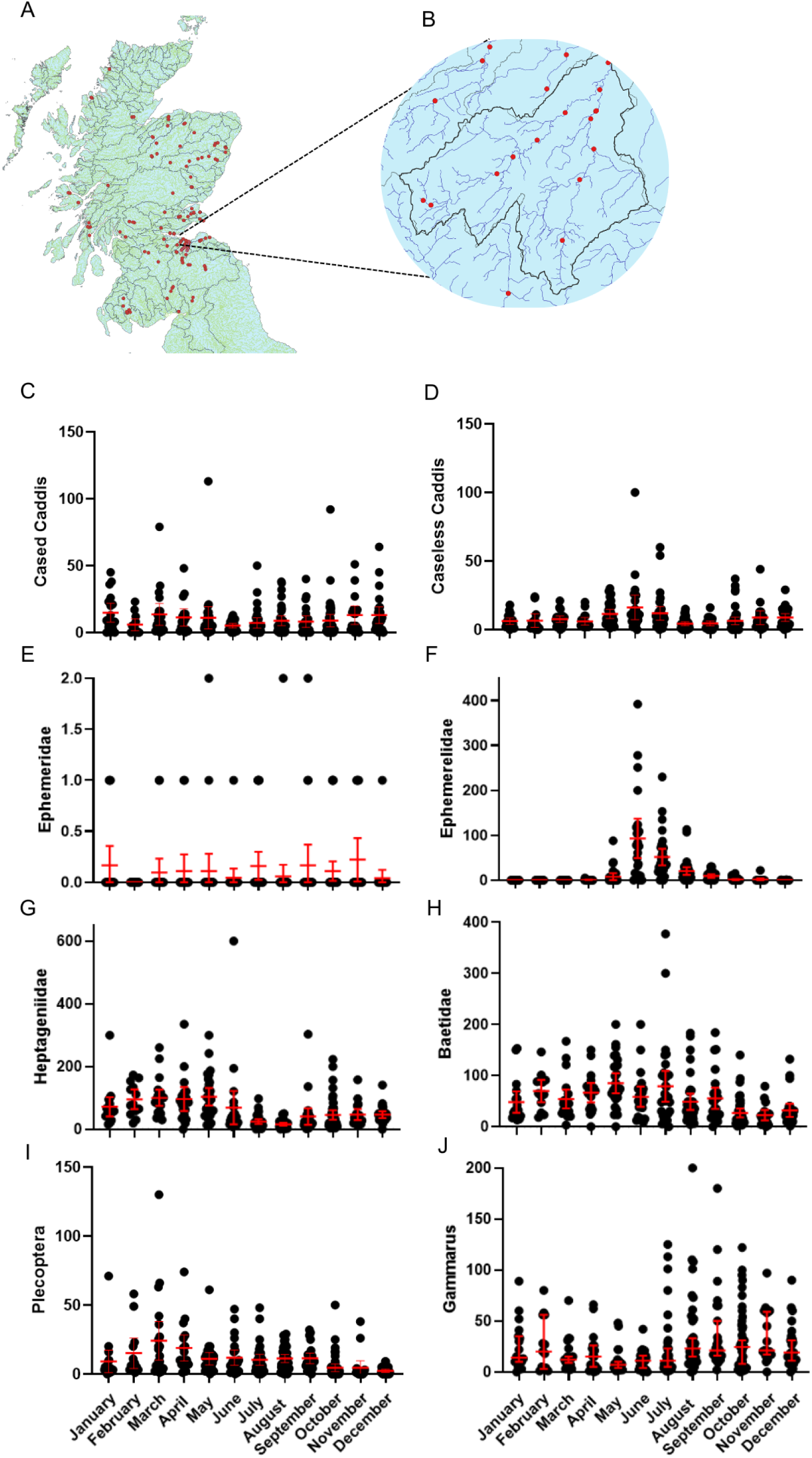
Seasonal variances in abundance of RMI taxa on the River Esk. A) Map of Scotland showing GooR sampling sites B) Expanded view of 13 study sites sampled by the “Riverfly-on-the-Esk’ GooR group (grey border defines catchment). C-J) Scatter plots showing number of invertebrates found in combined monthly data and mean +/-95% CI (indicated in red). Data collected from 2019 to 2025.

### Working with SEPA to establish trigger and target levels for RMI scoring

To set meaningful boundaries to data from GooR monitoring sites, and to allow volunteers and SEPA to work together to react to changes in monitoring site conditions, minimal acceptable (trigger) and optimal (target) levels were established for each GooR site. We used the Rivers Invertebrate Classification Tool (RICT) (46), combined with a SEPA derived conversion table to generate target, and trigger scores, respectively (see Materials and Methods for details). This method was then deployed to generate a national dataset of SEPA recognized targets and triggers for each of the 123 registered RMI monitoring sites.

There was a distinct separation between target and trigger values across the monitoring sites, with target values consistently clustering in the upper range of RMI scores (approximately 11–14) and trigger values clustering in lower RMI ranges (approximately 6–8) (Figure 4A). While clustering is evident, there is some variation within each band, more so in the targets which are less tightly clustered than the trigger levels, reflecting site specific differences. Although site-to-site variation is visible within each band, both distributions are relatively uniform, reflecting the consistent, and standardised threshold framework used to calculate the values across all monitoring locations.

**Figure 4.**
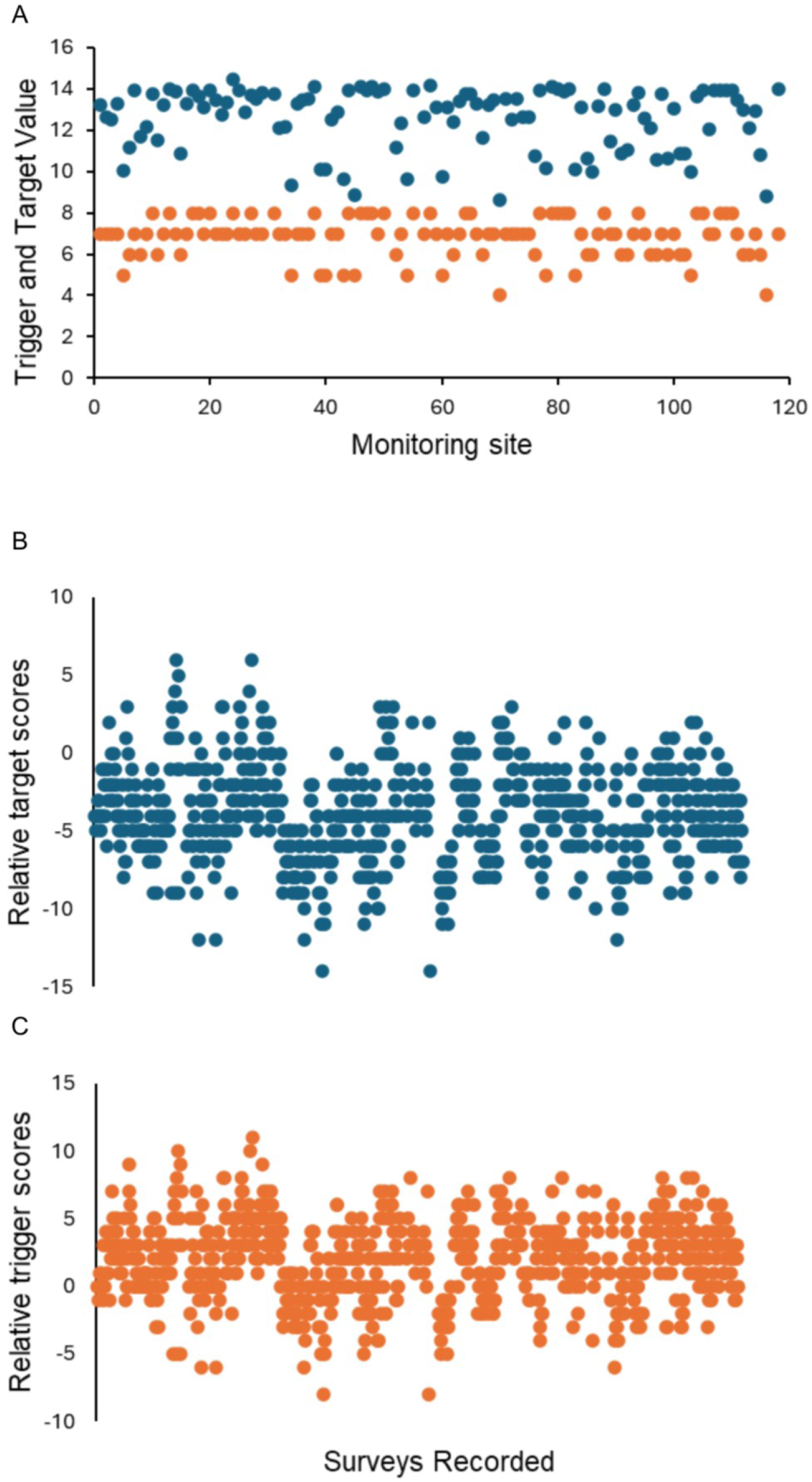
Establishing target and trigger scores for GooR monitoring sites (A) targets (blue) and triggers (orange) established for all 123 GooR sites B) Distribution of relative target scores (i.e. difference from target) across sampling sites from surveys in 2023-2025 (C) Distribution of relative trigger scores (i.e. difference from trigger) across sampling sites from surveys in 2023-2025.

The distribution of average RMI scores from each sampled site relative to the corresponding target level show a range between −14 and 6 (Figure 4B). A majority of sites lie below zero, indicating that most sites are not reaching target values. On the other hand, distribution of average scores from each sampled sites relative to corresponding trigger level show a range between −8 and 11. Most sites lie above zero, suggesting that a majority of sites are not breaching trigger levels.

**Figure 5.**
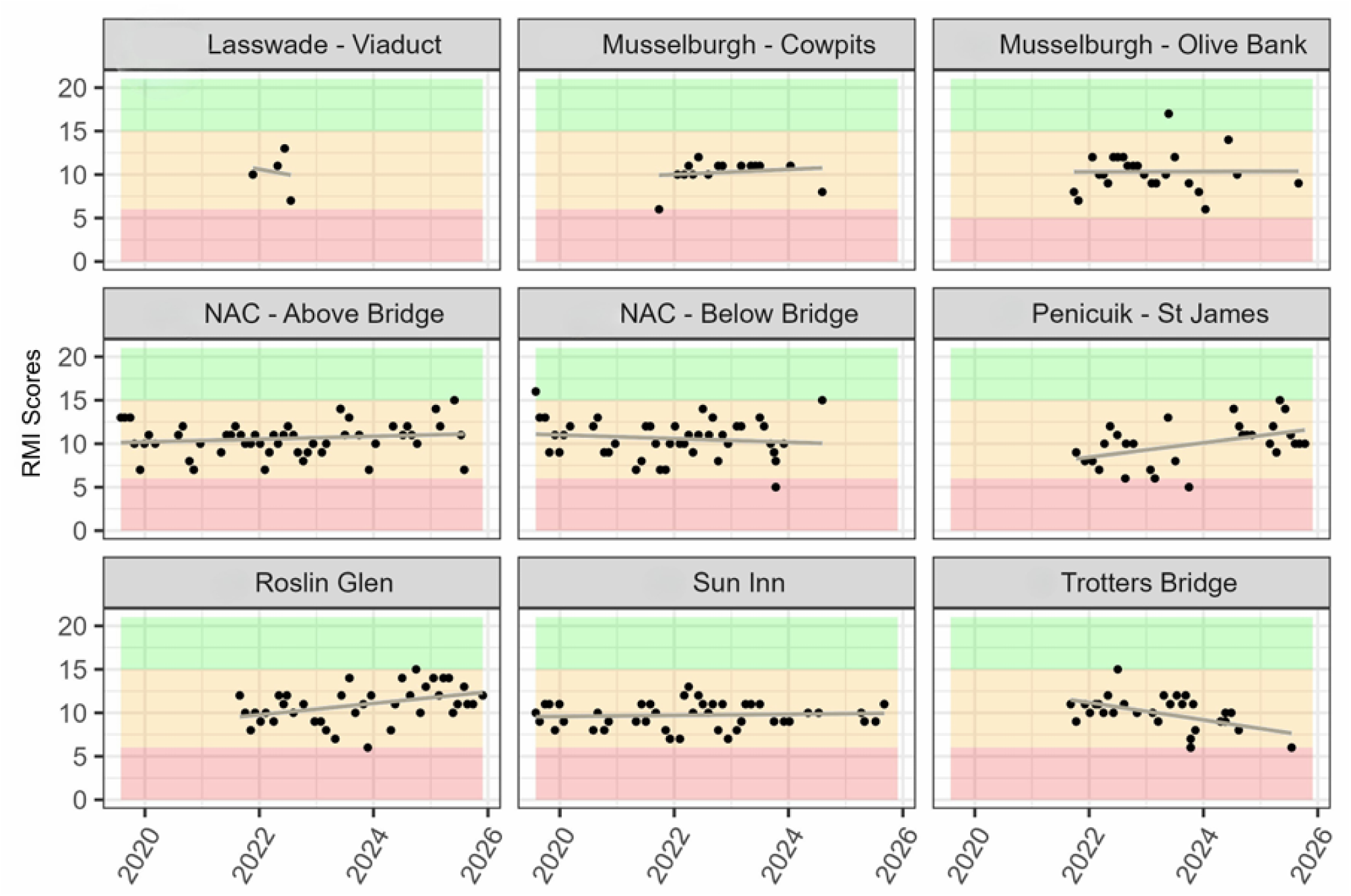
RMI scores over time at 9 monitoring sites on the Lothian Esk (site names in grey area). Green shaded area shows range above target, pink shaded area shows ranges below trigger levels, and yellow area represents an intermediate zone between the 2 thresholds. Trendline shows linear fit of data.

### Case study: Site specific trends on the Lothian Esk

Establishing target and trigger levels enabled GooR groups to quickly assess the health profile of a given site and monitor ‘breaches’ of trigger levels and ‘attainment’ of target levels. In some instances, groups began tracking multiple sites within a catchment. GooR Volunteers monitoring the Lothian Esk regularly monitored 9 sites along the river, each with distinctive features (Figure 5). Sites came on-line at different times in response to discussions with participants, and in some cases sampling was curtailed due to safety concerns (e.g. Lasswade Viaduct), or other logistical issues. From 2020-2025, near Trigger breaches were recorded at multiple sites over time (Figure 5, note data points near pink-yellow boundaries). Full trigger breaches were recorded at 2 sites (Figure 5, ‘NAC - Below Bridge’ and ‘Penicuik St James’).

**Figure 6.**
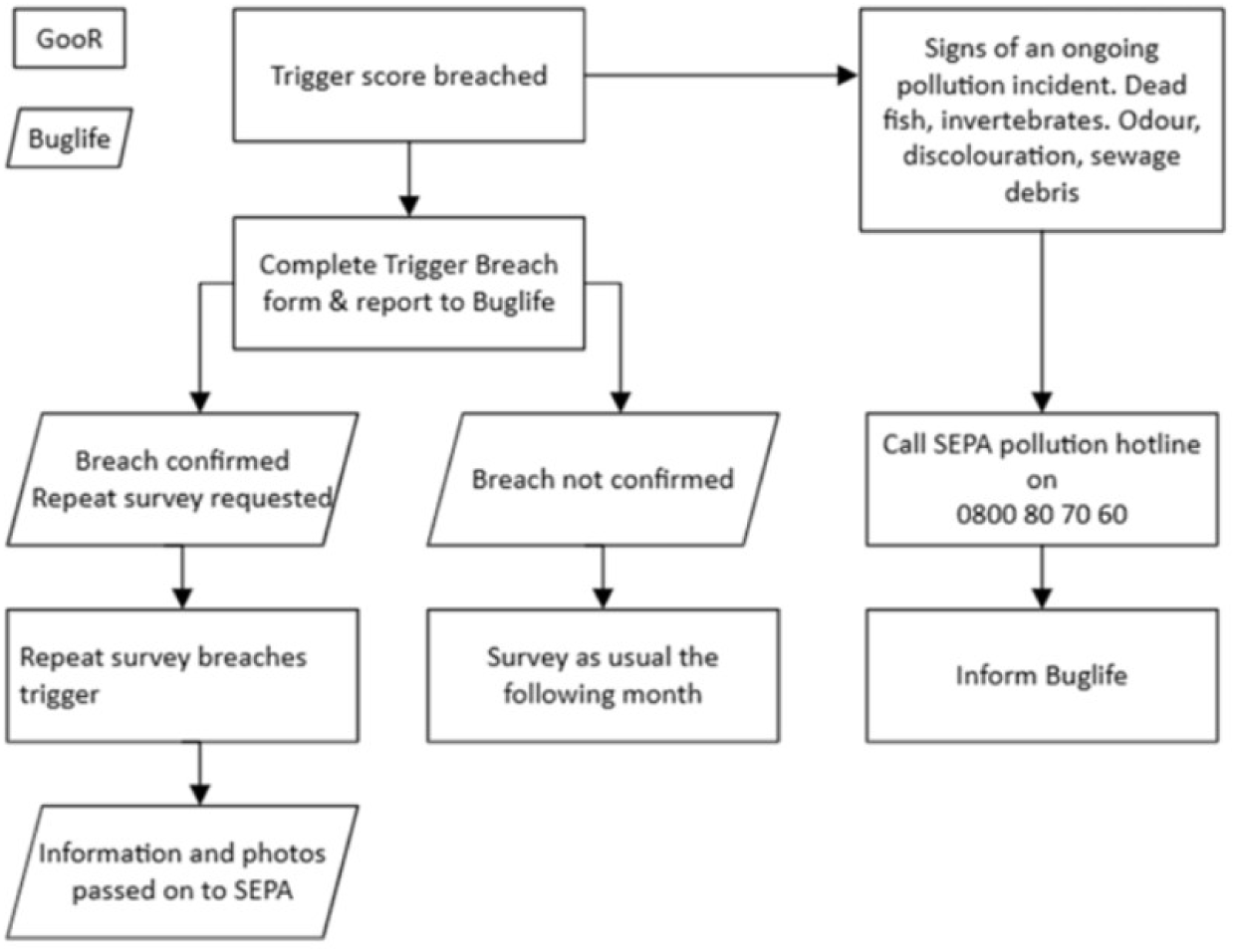
Procedure for reporting Trigger breaches to SEPA

Prior to the launch of GooR in October 2022, there was no formal working relationship between riverfly monitoring groups and SEPA. Data were collected but were not registered and/or integrated into SEPA operations and decision making. Bridging this divide was one of the major objectives of GooR. At the inception of GooR, Buglife personnel engaged with GooR groups and representatives from SEPA to iteratively co-develop a standardized procedure for reporting Trigger breaches (Figure 6). A decision tree, starting with a Trigger breach, provides clearly defined action pathways that lead to emergency hotlines that volunteers can call (in the case of obvious ongoing pollution events) or to further testing to confirm trigger breaches. A confirmed Trigger breach reported to SEPA then leads to a range of responses from written responses to site visits by SEPA ecologists. These processes and the working relationship between GooR and SEPA were formalized in 2025 in the form of a Memorandum of Understanding (MoU). Having this MoU in place ensures that trigger breaches are not just recorded but also formally presented to the relevant regulatory body. Breaches are sent from GooR volunteers to Buglife for verification, then on to SEPA for consideration and potential response.

**Figure 7.**
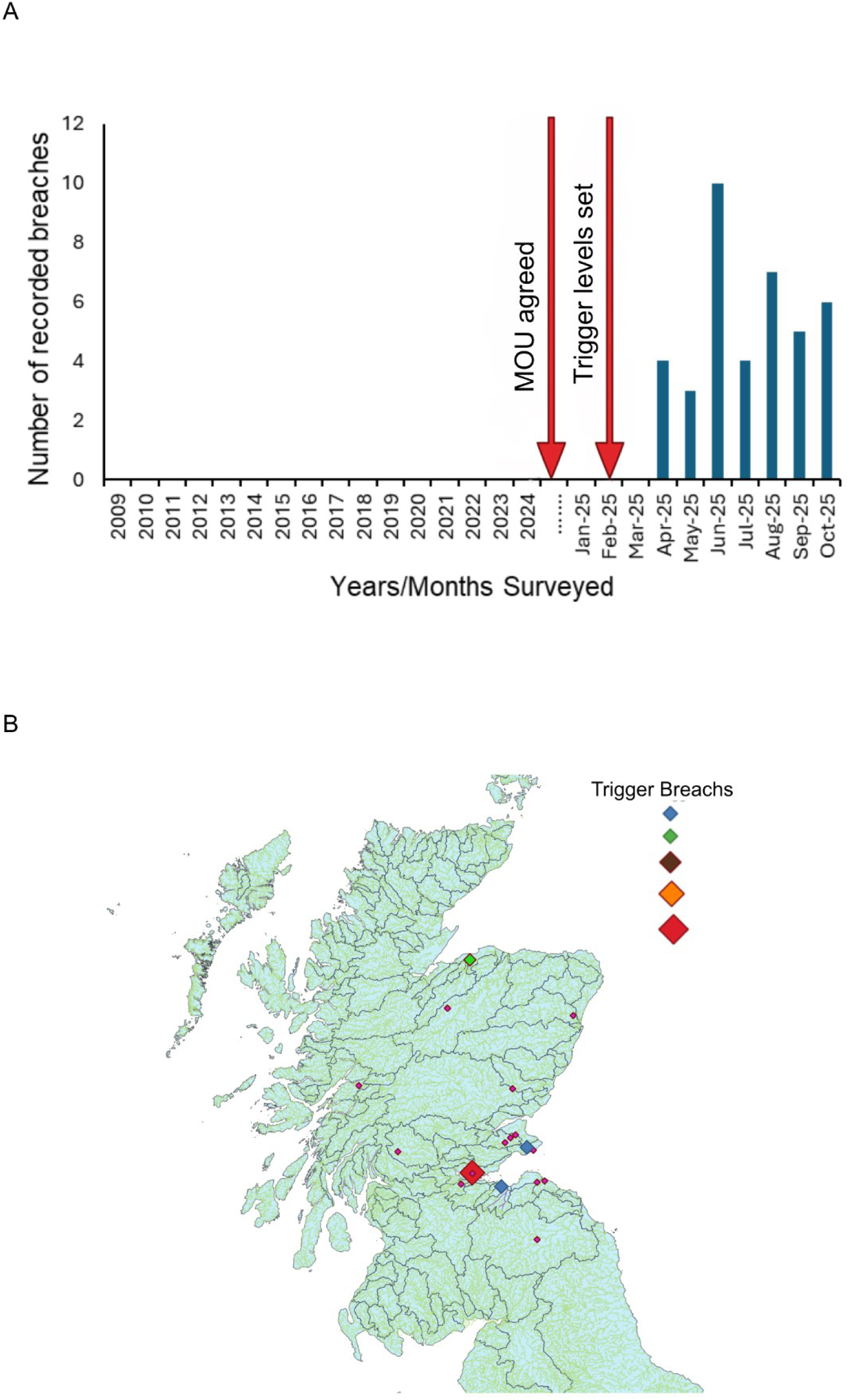
Formal relationship between Buglife, GooR and SEPA leads to registered, confirmed Trigger breaches. (A) Time points at which trigger breaches were reported directly to SEPA following the establishment of MoU and formal setting of the Trigger levels for all sites (B), Geographical distribution of Trigger breaches. Colour and size of diamond represents progression from single to multiple breaches

Across all GooR surveyed sites, 18 sites breached trigger levels from April to October 2025 (Figure 7A). Most Trigger breaches were single breaches, with 10 sites triggering a single time, followed by 2 Trigger breaches, recorded in 5 sites. The highest number of multiple Trigger breaches were recorded at sites around the Forth catchment with 5 breaches confirmed at 1 site, 4 in 2 others and 1 site with 3.

## Discussion

Citizen science networks that produce trusted, robust datasets over large areas have the potential to be transformative at local, national and international levels. Environmental Citizen Science projects range from terrestrial (47, 48, 49), freshwater (50) and marine habitats (51) and can involve a spectrum of engagement activities with varying levels of sophistication. Citizen scientists have revealed trends in population diversity, reduced pollution, and have helped to build cases for policy change (52).

### GooR as a model for growing a citizen science network

Freshwater citizen science projects such as Guardians of our Rivers (GooR), have the potential to play pivotal roles in determining whether government directives for water quality are indeed being met ‘on the ground’ in communities. For example, The Water Framework Directive (WFD) sets national goals for water quality in Scotland. The 2027 River Basin Management Plan formulated by WFD targets has articulated a goal of 81% of Scotland’s water bodies achieving at least “good” ecological status by 2027, as well as providing evidence of progress to delivering on ‘building more resilient rivers and communities’, a global Sustainable Development Goal (45). GooR research suggests that GooR data help contribute to these goals on a national and global stage, while also showing how citizen science can give a more nuanced picture of biological communities on a local level. Importantly, GooR also shows how it is possible to revitalize, expand, and restore momentum to existing citizen science networks. Within a period of 3 years, GooR managed to surpass the rate per annum of any of the previous 16 years of efforts to build Riverfly monitoring communities in Scotland - generating 70% of the data collected previously in just under 20% of the time. By considering the method and execution of the training, and the specific support packages offered, volunteers were recruited from across Scotland, increasing the coverage far beyond the central belt of Scotland into more remote highlands and islands communities. Judicious choice of sampling sites and recruitment of adequate numbers of volunteers per group ensured that the burden and reliance on individuals was kept to a minimum. Careful collection and curation of the data also ensured maximum value, utility, and confidence and allowed examination of multiple variables.

Given there had been past engagement with Riverfly in Scotland, it was important to take some learning from previously engaged volunteer groups prior to developing GooR. Informal feedback suggested that a lack of local support was a limiting factor, along with waning capacity in local groups. Our recruitment and training process took this feedback into account, and enhanced support was included (See Methods and Materials). Training kits were also provided free-of-charge to each new group to remove any barrier for groups getting up and running following completion of training. Face-to-face refresher sessions in the proceeding year offered a confidence boosts to volunteers. Data verification through online guidance and picture identification checks also ensured accessible volunteer support and increased confidence in the data for both volunteers and SEPA. The frequency of surveys carried out since 2009 shows clear patterns, with a boost from the launch of the CRIMP project in 2013-2015 showing what local support can achieve. This was reflected in 2019 with the rise of another local catchment project, Riverfly-on-the-Esk. However, focus on an individual catchment and limited funding constrained growth across Scotland. With the exception of 2020, when Covid 19 restrictions curtailed outdoor activities, the volume of survey data appeared to have plateaued as a result of groups becoming inactive. Volunteer engagement numbers for GooR showed a cumulative increase in volunteer numbers in consecutive years following the launch of GooR in 2022. However, it is evident that the potential for GooR has not been fully reached, as each year, there is a waiting list for training. Funding constraints limited further expansion of GooR in the past and will continue to be a challenge going forward unless further resource is allocated.

### GooR groups catalyze new research directions

Freshwater invertebrates can be found in virtually all aquatic habitats - from tree-borne epiphytic bromeliads to lake hypolimnion, from intermittent spring creeks to the brackish estuaries of great rivers. Studying these animals provides deep insights into the health of freshwater ecosystems (53). Use of aquatic invertebrates in professional research communities as bioindicators is widely accepted, with many national agencies and institutions having developed standardized biomonitoring protocols (54, 55) (56, 57, 58)). The degree to which citizen scientists can contribute novel observations to the scientific record (as opposed to simply reproducing known results or acting as technicians for professional researchers) is one measure of success of a citizen science program. Data collected by GooR volunteers over 3 years is indeed beginning to generate novel results. While the majority of RMI groups exhibited a fairly even distribution across the monitored sites, two groups (Ephemeridae and *Gammarus)* showed a strikingly high number of ‘absent’ recordings at sampling sites. This is in contrast to observations in England where both groups are relatively widely distributed (34) In Scotland, there is a single representative of the family Ephemeridae,(59) This species is associated with clean water(60) Historically, it had a highly restricted distribution in Scotland, probably as a result of historic river pollution, with small pockets located in headwater streams and unpolluted watercourses. More recently the distribution of this species has expanded; however, its exact range is unclear. GooR data provides foundational information on this potential expansion and provides a basis for exploring the differences in distribution in different regions of the UK. In contrast, there are four species of *Gammarus* known to be present (61). The most likely species to be encountered in running waters are *G. fossarum* and *G. pulex*, which are typically found in the lowland rivers. In Scotland, this limits their distribution to low lying areas in central, southern, and eastern Scotland. *Gammarus* as a genus are detrivores (62) which may be a limiting factor in their distribution; where riparian woodland is absent, their primary food sources may be more scarce. *Gammarus* are also intolerant of acidic conditions, so are typically absent from watercourses where local geological features reduce pH,(63) The new observations made by GooR groups about the distributions of RMI taxa now provide a foundation for further research into the reasons for observed geographical distributions. GooR observations also raise questions about whether the RMI metrics need to be rethought for use in Scottish waters.

GooR data also indicates precise locations of some RMI groups widely distributed across Scotland and this provides a basis for future work. Baetidae and Ephemerellidae distribution had the lowest percentage of ‘absent’ recorded from samples. Baetidae is a large family with mixed habitat preferences, with some species thought to being very common (e.g. *B. rhodani*) and others predicted to be particularly scarce (e.g. *B. digitatus*) (59). *B. rhodani* is also particularly tolerant of organic pollution, which makes it one of the most common mayfly species in rural areas of Scotland (64). Similarly, there are two species of Ephemerellidae in Scotland. *Serratella ignita and Ephemerella notata. S. ignita* has a very widespread distribution, due to its association with instream vegetation, such as mosses (59, 65), whereas *E. notata* is not often observed. GooR data is sensitive enough to show some of the basic distributions of freshwater invertebrate groups, which can then be used to gain a more nuanced understanding of local pressures and/or habitat types of common species, and to launch investigations on rare species.

### Growing relationships with diverse stakeholders

Building trust across multiple stakeholders is vital to the success of any citizen science network aimed at monitoring watercourses (66). This is facilitated by having a shared vision, clear communication, and mutual benefit. The national range of triggers and targets created by GooR broadly aligns with existing SEPA watercourse classification data, which suggests a majority of monitored Scottish rivers fall into 2 main condition categories (*High* or *Good)* with a smaller number in *Moderate*, *Poor,* or *Bad* (67). Comparing average scores from sampling sites to trigger and target levels revealed that most sites scored below the target score, suggesting that they are functioning below optimum conditions. For triggers, the majority of site scores lie above the trigger level; however, a number of sites had scores that were below the minimal acceptable conditions for their respective site. Initially there was no established response option in place for the majority of these events, so they went undocumented by SEPA. The formal partnership with SEPA in Scotland took several years to establish and was based on GooR building a strong network of trained volunteers able to generate robust multi-year datasets, which then could provide value to a formal relationship. We put in place a verification process for GooR data, along with a clear working arrangement. This helped ensure direct flow of robust data to SEPA as well as feedback to volunteers and, by extension, resulted in increased knowledge and understanding of previously unmonitored sites. Integral to this process was the establishment of trigger levels for all monitoring sites, providing a mechanism to detect decline in river health and initiate an appropriate response. While it is reassuring that, following the formal establishment of trigger levels in February 2025, only 39 triggers were breached, our analysis of the full GooR dataset (Figure 4) suggests that before that, trigger breaches would have gone undocumented, as the majority of GooR sites had no formal monitoring of breaches in place. One important aspect of the GooR-SEPA relationship is to acknowledge that some sites were known to have longstanding issues that would result in trigger breaches. These sites often required larger landscape scale solutions that were not immediately deliverable. Repeated trigger breaches at these GooR sites did not all lead to immediate interventions in the form of site visits by SEPA ecologists but were flagged as sites to watch for any further decline in quality, warranting further investigation. Open communication and sharing of local knowledge provide a foundation and focus for further discussion, investigations, and engagement across GooR groups, Buglife, academic researchers, and SEPA.

### Conclusions and outlook for future work

Overall, our sustained collaborative approach engaged the public in research, generated novel datasets, and reconfigured how a statutory body interacts with the public. Given the state of the environment and regular reporting of the degraded state of UK rivers during the launch and duration of the project (68, 69), we expected strong engagement. The public response to GooR exceeded expectations, leading to rapid and continuing growth of the network. GooR is made possible by funding from an array of sources across both the public and private sectors. The continued operation and expansion of citizen science efforts such as GooR hinge critically on continued support and funding from diverse sources. In particular, funding sources that support continuation of successful programs with demonstrated track records, as well as development of new ideas and approaches are critical. Ultimately, citizen science networks such as GooR provide value to people, governments and the environment, but sustaining success requires sustained collaboration and commitment from multiple partners.

## Materials and Methods

The framework for the method used in this research is based on the Riverfly Partnership’s Anglers’ Riverfly Monitoring Initiative (ARMI) (34). The Riverfly Partnership is a coalition of organisations and individuals from across the UK, including anglers, conservationists, scientists, water course managers and regulatory authorities, all working together to protect and improve the health and quality of rivers, and conserve riverflies. Now referred to as RMI, this method, launched nationally in March 2007, was developed in 2004 in response to rising public concern about the declining health of UK rivers and perceptions that regulatory authorities were not adequately detecting and remediating this decline due to limitations in resources and funding. In England where the technique was launched, training of volunteers and operation of the scheme is financed via matched funding; RMI groups raise money locally, including from local government, water companies and conservation trusts, as well as National Lottery grants, which is matched by the EA (Environment Agency 2019). Riverfly monitoring continues with 260 active groups monitoring rivers across England, collecting over 70,000 Samples to add to the dataset (34).

In Scotland, the first Riverfly group formed in 2009 on the River Avon, Falkirk, (training was also delivered to the River Tweed), but it wasn’t until the launch of the Clyde Riverfly Monitoring Partnership (CRIMP) project in 2013 that the number of sites began to grow in earnest (70). This project funded and supported the training of groups in the Clyde River Catchment. The Scottish Environmental Protection Agency dedicated a SEPA employee part-time to support the monitoring in Scotland. At its peak, there were 21 groups registered on the Riverfly database in Scotland, mainly focused around one river catchment. When funding for the CRIMP project ended and the SEPA officer retired, there was a decrease in engagement. By 2022 only 5 groups were still active, but sampling infrequently. Few groups had trigger levels (sampling score set at the minimal acceptable conditions for the site) set and there was no working relationship established with SEPA.

### Riverfly Monitoring Protocol

The RMI protocol is a simplified version of the Biological Monitoring Working Party scoring system for pollution monitoring (BMWP) (71), a biotic index widely used in the UK (72). It works on the premise that different taxa have different levels of sensitivity/ tolerance to organic pollution (73), and therefore the presence and abundance of particular taxa reflects distinct pollution levels. These taxa were chosen due to their expected distribution in rivers around the UK (particularly in England), year-round presence (except for Ephemerellidae), and because they are familiar to most anglers and easy to identify without specialized equipment (74). Presence and abundance of the larval stage of eight invertebrate groups is recorded so that major changes in water quality can be identified.

The eight “target groups” of invertebrates used in RMI are: Cased Caddis (Trichoptera); Caseless Caddis (Trichoptera); Mayfly (Ephemeroptera); Blue-winged Olive (Ephemerellidae); Flat-bodied (Heptageniidae); Olives (Baetidae); Stoneflies (Plecoptera); Freshwater Amphipods (*Gammarus spp.*) (Supplementary Figure 1). The collection method used for RMI is kick sampling, which is a common semi-quantitative method for sampling invertebrates in rivers. Invertebrates are sampled over a three-minute period, supplemented by a one-minute hand search of larger substrate clasts. Sampled macroinvertebrates are identified to families or orders (as appropriate), and assigned scores based on their abundances. This is synonymous with how macroinvertebrate-based ecological river health is derived for Water Framework Directive classifications, but at a coarser taxonomic resolution (75, 76). Sampling data is entered onto a public database hosted by the Freshwater Biological Association (34).

**Supplementary Figure 1.**
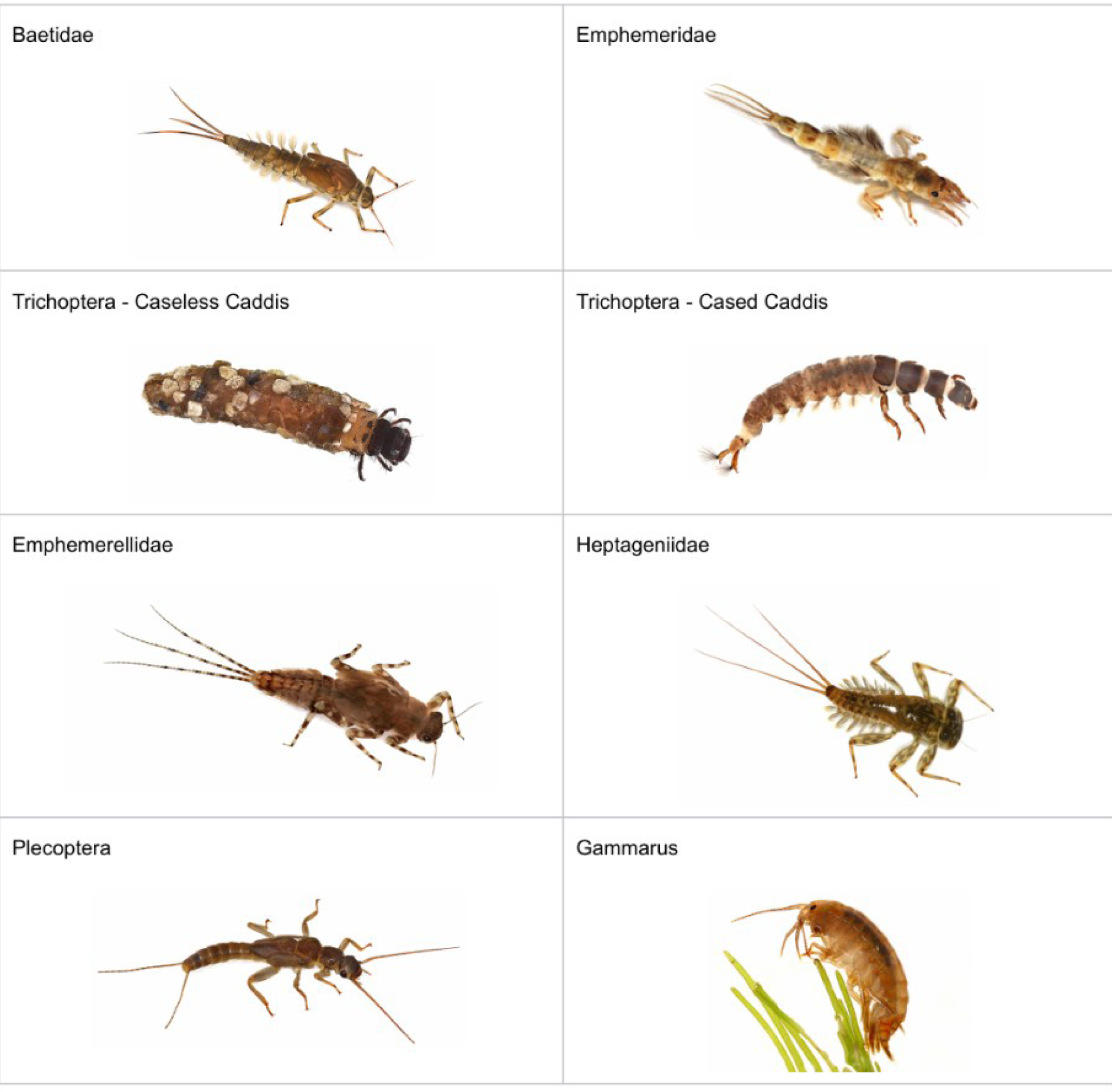
Representative examples of the 8 RMI taxa. Photos by Dr. Cyril Bennett MBE

### Volunteer recruitment and training

To engage communities in the project we chose the name ‘Guardians of our Rivers’. The aim was to instil a sense of active stewardship and togetherness for a joint cause. A press release was issued (October 2022) through multiple media channels across Scotland and using local networks to advertise the project and recruit volunteers. Volunteers contacted Buglife via email and phone to express an interest in the project. The flowchart below (Figure 8) shows the pathway used to induct volunteers:

**Figure 8.**
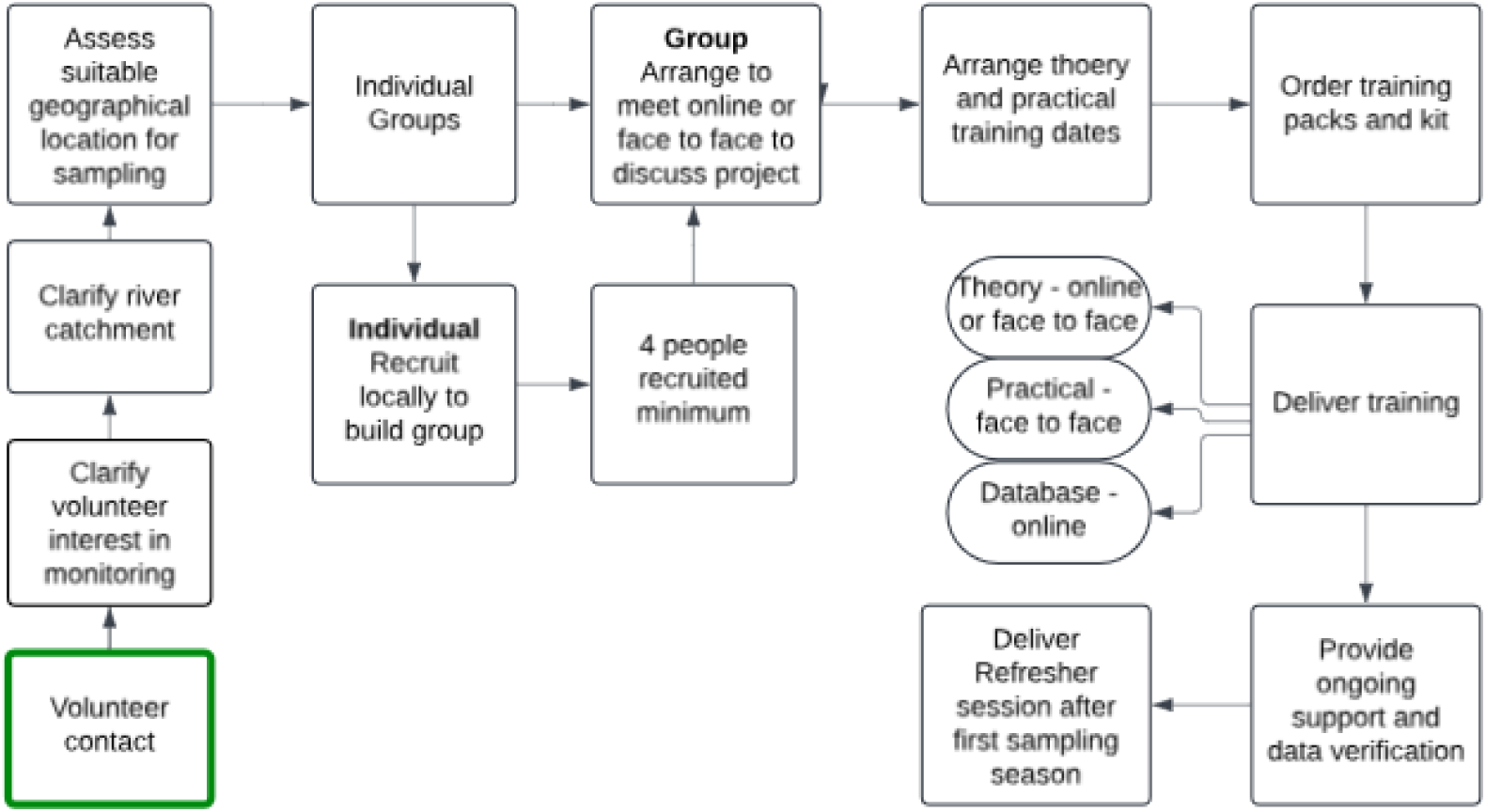
Flow chart describing GooR recruitment and training process.

Volunteer suitability for the project was discussed during this process with options given for volunteers to participate in multiple ways to suit both their personal and project requirements, with priority given to health and safety and practical sustainability.

### Monitoring site location

Monitoring locations were discussed during initial conversations with volunteers to take into account their local knowledge, needs and concerns. Consideration was also given to volunteer location within the river catchment to ensure that site location and access was sustainable for monthly monitoring. A desktop study of each site was undertaken to assess site access, geographical location and safety. The surrounding land use was checked for any potential pollution sources, modifications to the channel or significant barriers, water treatment plants or hydro-stations that would potentially influence the results. Permission to access sites was obtained if required. Monitoring site locations were confirmed following an onsite check and site risk assessment.

### Training packs

Each volunteer was provided with a training pack which included a laminated fold-out sheet, produced by the Field Studies Council (FSC), called the ‘Riverfly Monitoring Guide’. This fold-out gives an identification guide to the eight taxa of interest (specifically, high resolution images labelled with morphological identification features of interest), guidance on the sampling procedure and scoring system, advice on health and safety, and biosecurity, and instructions on how to report their scores. In addition to this, the pack contained a dynamic and generic risk assessment, biosecurity procedure and health and safety advice on Lyme’s disease and leptospirosis, and data recording sheets. Follow up supplementary information on invasive non-native species, sewage fungus, wildlife recording and spotting signs of pollution was provided digitally.

### Theory training

The theory section of the training was offered face-to-face or online. The online session was a 3-hour block delivered via cloud-based video conferencing platform Zoom. This standard presentation introduced the basis of biotic water quality monitoring and the RMI, explained the Riverfly Monitoring Guide and gave further identification advice on the eight target taxa, including videos showing their movement. Health and safety and biosecurity were also covered. We also demonstrated of this technique by presenting case studies from groups across the UK. These sessions were interactive with regular breaks so volunteers could meet each other as well as ask questions.

### Practical Training

Practical training was delivered onsite at the proposed monitoring location. Upon arrival, a site-specific risk assessment was completed, followed by a health and safety briefing covering in-stream sample collection and equipment use. Volunteers were shown the survey kit and observed an instream sample collection. They were then given time to take an individual sample, sort, identify, and record the eight RMI groups in the data sheet. On completion of both the theory and practical training, volunteers who were interested in carrying out monitoring were given a certificate.

### Equipment

Each group was given a full sampling kit to allow them to begin sampling immediately following completion of their training and to make sure that affordability wasn’t a barrier to engagement. Standardisation of both training and kit provided reduced the variability in the data collected and ensured that data could be communicated and discussed with SEPA more easily. The kit contained waders, gauntlet gloves, buoyancy aids, a sampling net (250-mm frame and 500-mm deep net bag with 1-mm mesh), bucket, dividing tray, sorting tray, sorting underlay mat, spoons, magnifying lenses and a large pipette.

### Sampling Method

At each selected monitoring site, volunteers identified the different habitat types present in the watercourse. Once identified, in stream sampling was divided proportionally according to the relative habitat areas. Volunteers use a standard sampling net to sample each site by taking a 3-min kick sample from the range of habitats present along a length of the site (length may vary depending on the characteristics of the water course). The volunteer then spends 1-min selecting large stones from the riverbed in the same stretch of the river, then hand washing them in the mouth of the net to dislodge organisms which may not have been collected in the kick sample. The volunteer subsequently cleaned the net contents of fine silt, emptied the remaining contents into a sorting tray from which scoring taxa were picked out live and sorted according to taxon in the segmented tray. The abundance of individuals within each RMI group was counted/estimated and awarded a score. The score for each taxon was summed to give an overall score for the sample, with higher scores indicating better water quality. A logarithmic scoring system was used for each taxon for each sample: Abundance Score Estimated Number A =1-9: B = 10-99: C=100-999: D= 100 1000+ (estimates the abundance of 8 macroinvertebrate target taxa on a logarithmic scale) (72)

### Health and Safety

To participate, every volunteer received full training including health and safety. Training packs included a copy of a generic and dynamic risk assessments were provided and a full site-specific risk assessment was undertaken at each site location during the practical training. Volunteers were asked to assess their general health prior to every monthly survey to check if they were fit to enter the water course. Volunteers who were not able or confident to enter the water course were invited to participate in the bank side sorting only. During training a first aid kit and hand sanitiser were provided.

### Follow up support and data verification

Following training, a package of support was offered to all participants. This included remote assistance through multiple channels; face to face refresher sessions; annual online gatherings and recruitment assistance. Survey data verification was undertaken through photographs sent from individual volunteers as well as data checks/approval through the database. Links to videos of each of the RMI 8 groups were shared to assist with identification.

### Triggers and Target setting

A key aspect of the method was to establish trigger and target levels for all sites on all rivers monitored by citizen scientists. Trigger levels are used to detect when significant changes in river health occur, which can be used to instigate investigations or more frequent monitoring by SEPA. The trigger level reflects the minimal acceptable conditions for the monitoring site and is calculated using Rivers invertebrate Classification Tool (RICT) software (77). RICT implements the River Invertebrate Prediction and Classification System (RIVPACS) model to determine the expected invertebrate community of a site based on landcover, hydrologic and geologic metrics, and alkalinity. The comparison between the invertebrate fauna expected in the absence of environmental stress and the fauna present provides a basis for assessing whether there has been a loss of ecological quality at a site. The ratio of the observed to predicted values of the biotic indices can be expressed as a series of Ecological Quality Indices (EQI) which can be used to define classes in a hierarchical manner (EQI bands)

### Data handling and analysis

Anonymized survey data were compiled, organized and plotted in QGIS, Microsoft Excel, Graphpad Prism, and R. Figures were prepared in Canva.

### Data availability

Underpinning data and analysis code are freely available upon request and online at: https://doi.org/10.17630/a17c9a84-aa85-443c-852b-22348716bd61

## Acknowledgements

This study would not have been possible without the support of the Riverfly volunteers across Scotland who gave their time to complete training, conduct surveys and enter data. Thanks to all members present and past of Riverfly-on-the-Esk, Riverfly on the Don, CLP Riverfly Group, Riverfly on the Dee, Aultbea Riverfly, Assynt Riverfly, Isle of Mull Riverfly, Falls of Clyde Riverfly, Annan Harbour Riverfly, River Avon Riverfly (Falkirk), River Deveron Riverfly, Kirrimuir Rifflers, Kinnesburn Riverfly, Dreel Burn Riverfly, Kinlochbervie Riverfly, River Spey Riverfly, River Findhorn Riverfly, Cairngorm Rangers Riverfly, Balmoral Riverfly, Inch Marshes Riverfly, Durell at Dalnacardoch Riverfly, Lochgilphead Riverfly, Annon Harbour Riverfy, Mar Lodge Riverfly, Guardians of the Tweed, Figgate Park Riverfly, River Eden and Motray Riverfly, River Peffery Riverfly, Breich Water Riverfly, Glencoe Riverfly, River Cree Riverfly, River Add Riverfly, River Esk (Dunfries and Galloway) Riverlfy, Water of Ae Riverfly, Water of Chon Riverlfy, Action West Loch Tarbet, River Nith, Loch Lomond, Lunan Burn, R. Avon (S. Lanarkshire), River Annan Catchment Riverfly and Tyne Catchment Riverfly.

Thanks also to the Riverfly Partnership and Cartographer for data sharing, with special thanks to Steve Brooks and John Clayton for supporting training. Thanks to the team at Buglife for the support and The Clyde River Foundation for initial studies. We also thank SEPA for partnering with us to establish target, triggers and response pathways.

Special thanks to Mica Lewis-Biddick.

## Grants

This work was supported by Buglife Scotland with grants from NatureScot, Swire Charitable Trust, Animal Friends Pet Insurance, Hugh Fraser Foundation, The Northwick Trust, National Lottery Heritage Fund, Mackies, and Highlands and Islands Environment Fund, Cairngorms Trust and support from the Riverfly Partnership, Midlothian Council, Cala homes, Midlothian Climate Action Hub, and Musselburgh District Angling Association. This work was also supported by a PhD studentship from the Sir Harold Mitchell Fund (University of St Andrews).

## Disclosures

No conflicts of interest, financial or otherwise, are declared by the authors.

## Notes

https://doi.org/10.17630/a17c9a84-aa85-443c-852b-22348716bd61

### Competing Interest Statement

The authors have declared no competing interest.

### Summary of Updates

Update to authors name on manuscript and figure.

